# Porcn is essential for growth and invagination of the mammalian optic cup

**DOI:** 10.1101/2022.09.01.506058

**Authors:** Sabine Fuhrmann, Sara Ramirez, Mirna Mina Abouda, Clorissa D. Campbell

**Affiliations:** Dept. of Ophthalmology and Visual Sciences; Vanderbilt Eye Institute, Vanderbilt University Medical Center; Dept. of Cell and Developmental Biology, Vanderbilt University Medical School; Nashville, TN

**Keywords:** microphthalmia, focal dermal hypoplasia, Goltz Syndrome, Porcn, Wnt, optic vesicle, optic cup, mouse, embryo

## Abstract

Microphthalmia, anophthalmia, and coloboma (MAC) are congenital ocular malformations causing 25% of childhood blindness. The X-linked disorder Focal Dermal Hypoplasia is frequently associated with MAC and results from mutations in *Porcn*, a membrane bound O-acyl transferase required for palmitoylation of Wnts to activate multiple Wnt-dependent pathways. Wnt/β-catenin signaling is suppressed in the anterior neural plate for initiation of eye formation and is subsequently required during differentiation of the retinal pigment epithelium (RPE). Non-canonical Wnts are critical for early eye formation in frog and zebrafish, however, it is unclear whether this also applies to mammals. We performed ubiquitous conditional inactivation of *Porcn* in mouse around the eye field stage. In *Porcn*^*CKO*^, optic vesicles (OV) arrest in growth and fail to form an optic cup. Ventral proliferation is significantly decreased in the mutant OV, with a concomitant increase in apoptotic cell death. While pan-ocular transcription factors such as PAX6, SIX3, LHX2, PAX2 are present, indicative of maintenance of OV identity, regional expression of VSX2, MITF, OTX2 and NR2F2 is downregulated. Failure of RPE differentiation in *Porcn*^*CKO*^ is consistent with downregulation of the Wnt/β-catenin effector LEF1, starting around 2.5 days after inactivation. This suggests that *Porcn* inactivation affects signaling later than a potential requirement for Wnts to promote eye field formation. Altogether, our data shows a novel requirement for Porcn in regulating growth and morphogenesis of the OV, likely by controlling proliferation and survival. In FDH patients with ocular manifestations, growth deficiency during early ocular morphogenesis may be the underlying cause for microphthalmia.

## Introduction

Congenital ocular malformations – <u>m</u>icrophthalmia (small eye), <u>a</u>nophthalmia (absent eye, also called extreme microphthalmia), <u>c</u>oloboma (optic fissure closure defect in the ventral OC) (collectively hereafter MAC) – originate from defective morphogenesis during early eye development and cause 25% of childhood blindness (Clementi et al., 1992; Graw, 2019; Morrison et al., 2002). Eye development is initiated during gastrulation in a single domain in the anterior neural plate, the eye field, characterized by expression of eye field transcription factors (EFTFs) that include Pax6, Six3, Six6, Rx, Tbx3 and Lhx2 (Bailey et al., 2004; Liu and Cvekl, 2017; Motahari et al., 2016; Oliver et al., 1995; Zuber et al., 2003). In combination with secreted factors inhibiting Nodal, BMP and Wnt/β-catenin signaling, EFTFs form a gene regulatory network to promote eye field specification (Zuber et al., 2003). Subsequentially, the eye field separates into two optic pits that evaginate laterally. Bilateral optic vesicles (OVs) are formed, comprised of neuroepithelial progenitor cells giving rise to optic stalk, retina and RPE. The distal domain of the OV contacts the overlying surface (lens) ectoderm, and both OV and ectoderm invaginate to form the optic cup (OC) and the lens vesicle, respectively. While the coordinated interactions regulating regionalization of OC and OV are well studied (for reviews, see (Fuhrmann, 2010; Miesfeld and Brown, 2019; Viczian, 2014)), regulation of the morphogenetic events during OV evagination and OC invagination are less understood.

Focal Dermal Hypoplasia (FDH; Goltz Syndrome) is an x-linked dominant disorder resulting from abnormal mesodermal and ectodermal development and is frequently associated with MAC (Gisseman and Herce, 2016; Goltz et al., 1962; Temple et al., 1990; Wang et al., 2014). FDH is caused by mutations in *Porcn*, a membrane bound O-acyl transferase localized to the endoplasmic reticulum (Grzeschik et al., 2007; Wang et al., 2007). Porcn is mostly dedicated to palmitoylate the Wnt family of cysteine-rich secreted glycoproteins, and this modification is necessary for trafficking and signaling (Galli et al., 2007; Tanaka et al., 2000). Porcn is expressed in the developing mouse eye (Biechele et al., 2011). We have shown that conditional *Porcn* disruption at the OV stage results in coloboma and RPE defects in most embryos, among other abnormalities (Bankhead et al., 2015).

Wnt ligands bind to surface receptors, including Frizzled transmembrane receptors (Fzd), and activate canonical (Wnt/β-catenin) and non-canonical pathways. Wnt/β-catenin activation prevents cytoplasmic degradation of β-catenin, thereby allowing its nuclear translocation to activate TCF/LEF transcription factors. Wnt/β-catenin exerts many distinct functions during different phases of eye development (for reviews, see (Fuhrmann, 2008; Fujimura, 2016)). During early OC morphogenesis, it is required for RPE differentiation (Fujimura et al., 2009; Hagglund et al., 2013; Westenskow et al., 2009) and dorsoventral patterning (Veien et al., 2008; Zhou et al., 2008). The pathway is tightly regulated; ectopic activation by disruption of the antagonists Dkk1 and Axin2 can result in microphthalmia and coloboma (Alldredge and Fuhrmann, 2016; Lieven and Ruther, 2011). During forebrain development, multiple antagonists maintain anterior neural fate. Hyperactive Wnt/β-catenin frequently leads to posterization and rostral truncation of the forebrain, with concomitant severe decrease of EFTF expression. The EFTF Six3 can directly suppress Wnt/β-catenin signaling (Lagutin et al., 2003), providing another mechanism for precise coordination of pathway activity.

Non-canonical, β-catenin-independent Wnt signaling occurs via multiple pathways, including intracellular Ca^2+^ release, activation of JNK and Planar Cell Polarity (PCP). It is essential for eye field formation in non-mammalian vertebrates by different mechanisms; it represses Wnt/β-catenin activity and promotes expression of EFTFs in frog and zebrafish (for review, see (Fuhrmann, 2008)). Close to the caudal border of the eye field, non-canonical Wnt11 and Wnt4 act through Fzd5 and Fzd3 to promote expression of the EFTFs Pax6 and Rx (Cavodeassi et al., 2005; Maurus et al., 2005; Rasmussen et al., 2001). In addition, Wnt11 and crosstalk between JNK/PCP and ephrinB1 are required for morphogenetic movements of ocular progenitor cells into the eye field (Cavodeassi et al., 2005; Cavodeassi et al., 2013; Heisenberg et al., 2000; Lee et al., 2006; Moore et al., 2004). However, it is unknown whether non-canonical Wnt signaling functions similarly in mammals; for example, germline disruption of Wnt4, Wnt11 and Fzd3 does not cause obvious ocular phenotypes, suggesting either redundancy of pathway components and/or context- and species-dependent mechanisms.

To investigate the role of Porcn before OV morphogenesis, we performed temporally controlled inactivation before and during the eye field stage when Wnts are expressed in the cranial region (Kemp et al., 2005; Kispert et al., 1996; Parr et al., 1993; Yamaguchi et al., 1999). Our results show that *Porcn* inactivation around eye field formation leads to severely arrested OV three days later. Key regulatory genes for RPE differentiation OTX2, MITF and NR2F2 are absent, proliferation and survival of ocular progenitors in the OV is decreased, and invagination during OC morphogenesis fails. Our studies reveal a novel role for Porcn during earliest stages of mouse eye development, recapitulating severe microphthalmia in FDH.

## Method

Mouse lines were maintained on a mixed genetic C57BL/6 and CD-1 background. For temporally controlled *Porcn* inactivation, a conditional *Porcn* allele, tamoxifen-inducible, ubiquitous *Gt(ROSA)26Sor*^*tm1(cre/ERT)Nat*^ (hereafter *ROSA26*^*CreERT*^) and the recombination reporter *RosaR26* were utilized, with established genotyping protocols or Transnetyx (Cordova, TN) using Taqman with custom-designed primers (Badea et al., 2003; Bankhead et al., 2015; Barrott et al., 2011; Soriano, 1999; Sun et al., 2020). Noon of the day with an observed vaginal plug was counted E0.5. Pregnant dams received tamoxifen (0.1 mg/g) by oral gavage between E6.5 and E7.5 (Park et al., 2008). To detect proliferating cells, pregnant dams received one intraperitoneal EdU injection two hours before sacrificing (30 µg/g; Thermofisher/Invitrogen; #E10187). Male embryos with conditional deletion of *Porcn* (hereafter *Porcn*^*CKO*^) and control littermates were processed as previously published (Sun et al., 2020). For antigen retrieval, cryostat sections were treated with 1% Triton X-100 or hot citrate buffer (pH 6). Detailed antibody information is provided in Suppl. Table 1. Filamentous actin was detected using Phalloidin (1:50; Thermo Fisher Scientific; #A12379). ApopTag Fluorescein In Situ Apoptosis Detection Kit (EMD Millipore, #S7110) was used to detect apoptotic cells. For EdU detection, the Click-iT® EdU Imaging Kit (Thermo Fisher Scientific; #C10637) was utilized. Cryostat sections were counter-labeled with DAPI and mounted in Prolong Gold Antifade. No developmental defects were observed in controls: conditional heterozygous female embryos (hereafter *Porcn*^*CHET)*^ and embryos with or without *Cre*. Unless otherwise indicated, a minimum of 3 embryos from at least 2 individual litters were analyzed per genotype, time point and marker. For analyses of E9.5 embryos induced at E6.5, 3 embryos were analyzed and 2 showed indication of ocular development.

Images were captured using a U-CMAD3 camera, mounted on an Olympus SZX12 stereomicroscope, and a XM10 camera, mounted on an Olympus BX51 epifluorescence system. For confocal imaging, the Olympus FV100 or ZEISS LSM 880 systems were used. Images were processed using ImageJ (NIH) and Adobe Photoshop software (versions CS6, 23.3.1).

Quantification of EdU-labeled cells was performed on cryostat sections midway through the OV, with separation of dorsal and ventral subdivisions. The percentage of cells incorporating EdU was calculated by determining the number of total cells using DAPI-labeled nuclei. Statistical analysis was performed using Prism version 9 (Graphpad) for Student’s T-test.

## Results

### Conditional Porcn inactivation before eye field induction (E6.5) causes abnormal OV morphogenesis

We observed that Porcn needs to be depleted in multiple ocular and extraocular tissues at the OV stage to recapitulate consistently the ocular abnormalities found in FDH patients (Bankhead et al., 2015). In mouse, the eye field becomes established at E7.5, and Wnts are expressed in tissues adjacent to the eye field. To target both cell autonomous and non-cell autonomous Wnt production, we disrupted *Porcn* using the ubiquitous *ROSA26*^*CreERT*^, allowing temporally controlled recombination. To account for delay of pathway inactivation in responding cells due to cell non-autonomous Wnt ligand production, extracellular release and downstream receptor activation, we started inactivating *Porcn* at E6.5. *RosaR26* recombination confirmed ubiquitous *Cre* activity, as expected (Suppl. Figure 1) (Soriano, 1999). At E8.5-8.75, *Porcn*^*CKO*^ show severe posterior truncation (Barrott et al., 2011; Biechele et al., 2013). However, anterior regions do continue to develop, with reduced size of OVs (8-13 somites; Figure 1B-H). In *Porcn*^*CKO*^, expression of the Wnt/β-catenin readout LEF1 can be slightly decreased in the dorsal OV, consistent with loss of Wnt production (Figure 1D). Lhx2 is required for the transition from eye field to OV and is robustly expressed in *Porcn*^*CKO*^ (Figure 1F). Otx2 is critical for rostral brain regionalization and early ocular development. In *Porcn*^*CKO*^, OTX2 expression is present in the dorsal OV (Figure 1H). To determine whether the observed alterations could be due to a developmental delay, we examined *Porcn*^*CKO*^ embryos 1-2 days later. However, we observed major developmental abnormalities in E10.5 *Porcn*^*CKO*^ embryos, not suitable for further analysis (not shown). Thus, we continued to examine embryos at E9.5 (20-23 somites); *Porcn*^*CKO*^ exhibit more defects compared to E8.5-8.75, including abnormalities in the head region (Figure 1J-K). Morphogenesis of the OVs does not proceed properly; they are arrested and fail to contact the surface ectoderm (Figure 1M, O, Q, S). Expression of LEF1 is absent in the dorsal OV, indicating loss of Wnt/β-catenin signaling (Figure 1M). Phalloidin labeling reveals that apicobasal polarity is generally preserved (Figure 1O), thus, it is possible that non-canonical Wnt pathways such as planar cell polarity are largely maintained. Robust LHX2 expression is maintained in *Porcn*^*CKO*^ (Figure 1Q). In contrast, OTX2 is downregulated in the dorsal OV, while weak expression is detectable in the adjacent forebrain (Figure 1S). Overall, our results demonstrate that *Porcn* inactivation starts to affect eye development after approx. 2.5 days, with slightly decreased Wnt/β-catenin signaling in the dorsal OV. Evagination can occur, suggesting that OV identity is maintained, and morphogenesis initiated. However, with ongoing loss of Porcn at E9.5, dorsal regionalization and expansion of the OV is severely affected.

**Figure 1:**
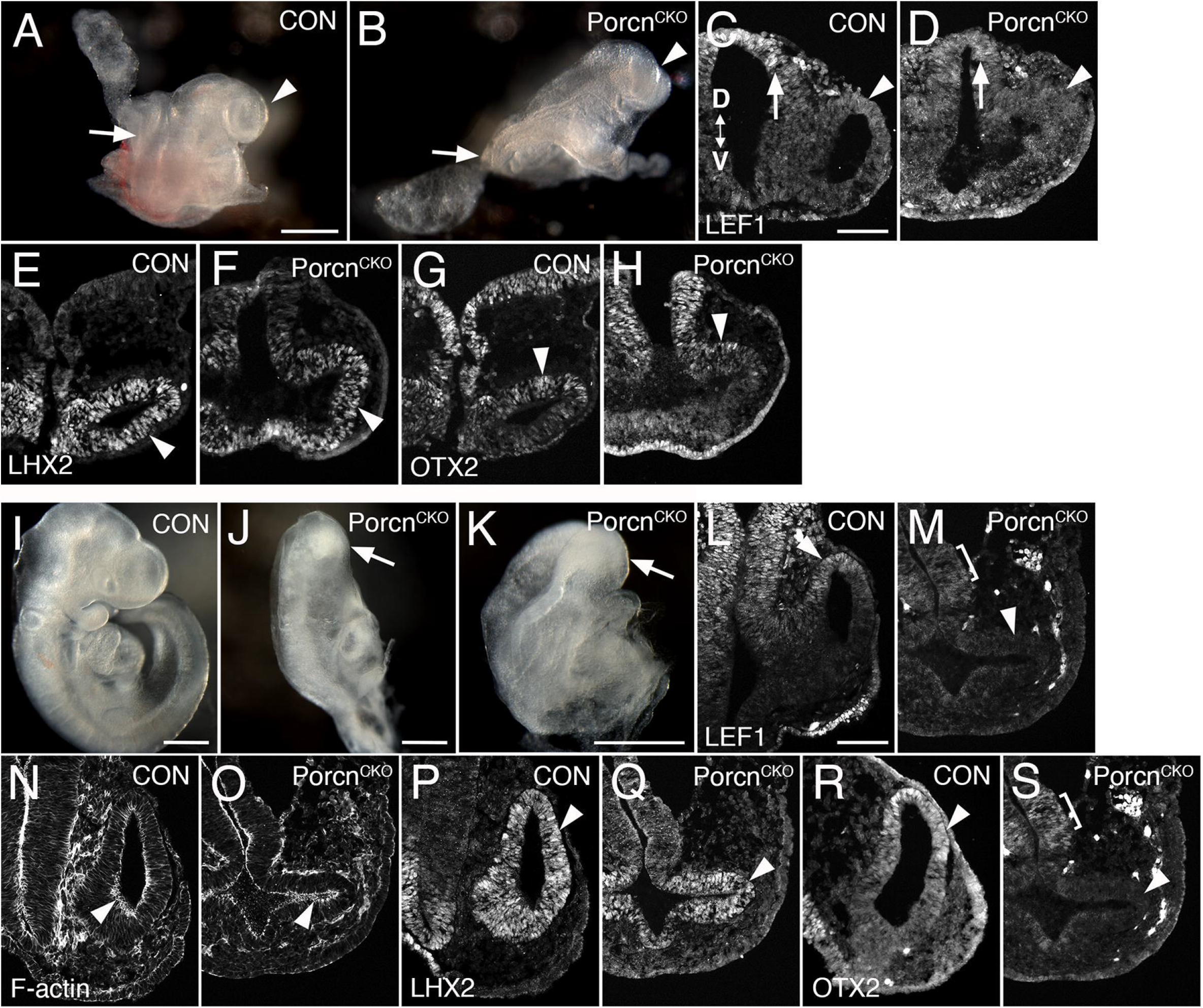
Conditional *Porcn* inactivation before eye field formation (E6.5) interferes with early OV morphogenesis. (A-H) Embryos at E8.75-E9.0. (A) Control embryo (8 somites) with OVs (arrowhead). (B) *Porcn*^*CKO*^ with caudal truncation (arrow; somites not detectable). Head development proceeds at this stage and OV-like structures are present (arrowhead). (C, D) LEF1 expression in 13 somite embryos. Coronal cryostat sections show slightly decreased LEF1 expression in the dorsal OV of *Porcn*^*CKO*^ (D; arrowhead), expression in the dorsal and ventral forebrain is largely maintained (D; arrow). E-H: 7-8 somite embryos. (F) LHX2 is robustly expressed in the mutant OV (arrowhead). (H) In *Porcn*^*CKO*^, OTX2 expression is present in the dorsal OV (arrowhead). (I-S) Embryos at E9.5 (20-23 somites); 2 of 3 embryos showed distinct OVs, shown here in J-S. (J-K) Examples of *Porcn*^*CKO*^ showing severe abnormalities throughout the body, including the heads (arrows). (M) LEF1 expression is absent in *Porcn*^*CKO*^ OV (arrowhead), low expression is detectable in the adjacent diencephalon (bracket). (O) Phalloidin labeling shows that apicobasal polarity is largely maintained in *Porcn*^*CKO*^ (arrowhead). (Q) In the mutant OV, LHX2 is robustly expressed, similar to controls (arrowhead). (S) OTX2 is absent in the OV of *Porcn*^*CKO*^ (arrowhead). Weak expression is detectable in the forebrain neuroepithelium (bracket). OV OV. D<->V: Dorsoventral orientation. Scale bars A, I-K: 0.5 mm; C, L: 0.1 mm.

### Porcn deletion during eye field induction (E7.5) variably affects brain and eye formation

To bypass major developmental defects before OV morphogenesis (Figure 1), we administered tamoxifen one day later (E7.4-7.5). At E10.5, *Porcn*^*CKO*^ embryos were recovered with a range of ocular abnormalities (Figure 2). In less affected *Porcn*^*CKO*^, telencephalic vesicles are slightly reduced (Figure 2F). Eye size can be reduced, associated with incomplete OC morphogenesis, failure of lens vesicle formation and accumulation of POM between distal OV and surface ectoderm (Figure 2G-J). In mildly affected *Porcn*^*CKO*^, OVs can be more closely associated with lens ectoderm, occasionally with further advanced invagination (not shown). We observed that tamoxifen treatment at E7.4 results in a higher proportion of *Porcn*^*CKO*^ embryos with a more severe ocular phenotype (Figure 2K-S). This is most likely due to developmental variability between litters and embryos at the time of recombination. More severely affected *Porcn*^*CKO*^ exhibit defects in mid- and forebrain development; the mid-hindbrain border is missing and telencephalic vesicles are underdeveloped (Figure 2K). Ocular morphogenesis is consistently arrested with small OVs that fail to expand, with abnormal accumulation of anterior POM preventing close contact between distal OV and lens ectoderm (Figure 2L-O). Consequently, neither the OV nor the surface ectoderm show any signs of invagination (Figure 2M-O). Some thickening of the surface ectoderm is occasionally detectable, suggesting that early aspects of lens morphogenesis can be initiated (Figure 2M). In *Porcn*^*CKO*^ embryos with a severe phenotype, overall LEF1 expression is decreased, specifically in the dorsal forebrain, dorsal OV and facial primordia, indicating widespread downregulation of the Wnt/β-catenin pathway (Figure 2Q, S).

**Figure 2:**
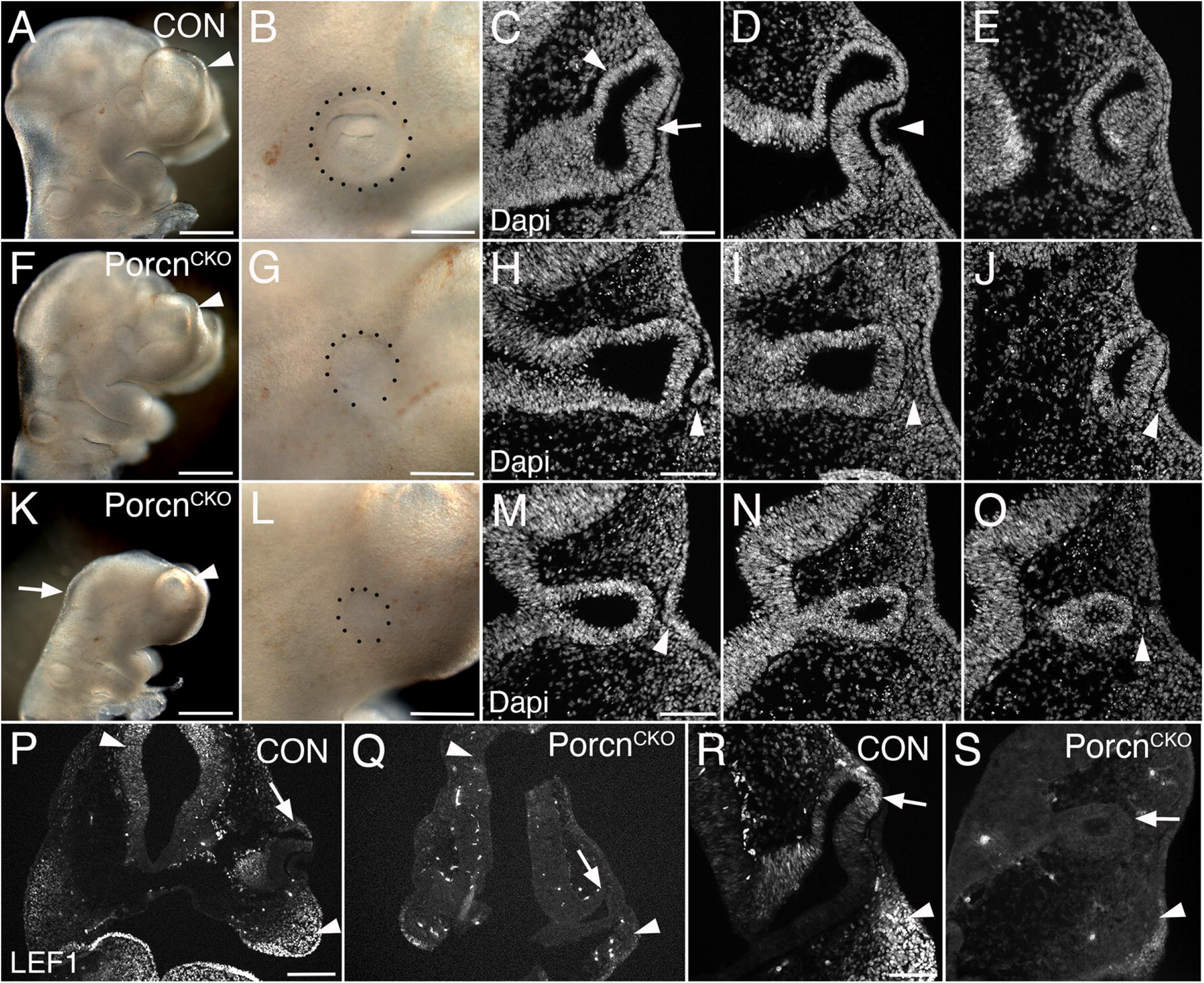
Conditional *Porcn* inactivation during eye field formation (E7.4-7.5) causes a range of brain and ocular abnormalities at E10.5. (A-E) Control *Porcn*^*HET*^ embryos showing telencephalic vesicle (A; arrowhead), early OC with lens vesicle (B; dotted outline), presumptive RPE and neural retina (C; arrowhead and arrow, respectively) and invaginating lens vesicle (D; arrowhead) in sequential Dapi-labeled coronal sections (C-E). (F-J) *Porcn*^*CKO*^ with mild abnormalities; reduced telencephalic vesicle size (F; arrowhead), reduced eye size (B), incomplete invagination of OV and lens ectoderm (H, J; arrowheads) and accumulation of POM between distal OV and lens ectoderm (I; arrowhead). (K-O) *Porcn*^*CKO*^ with major defects showing loss of the mid-hindbrain border and further reduction of telencephalic vesicles (K; arrow and arrowhead, respectively), microphthalmia (L; dotted outline, compare B, G, L), arrested OV (M-O) and accumulation of POM (O; arrowhead). Some thickening of the surface can occur (M; arrowhead). (P-S) LEF1 immunolabeling of coronal control sections shows expression in the dorsal forebrain (P, left arrowhead), facial primordia mesenchyme (P; right arrowhead, R; arrowhead) and dorsal OC (P, R; arrows). In *Porcn*^*CKO*^ embryos, LEF1 is severely reduced in the dorsal forebrain (Q; left arrowhead), facial primordia (Q; right arrowhead, S; arrowhead) and dorsal OC (Q, S; arrows). Scale bars A, F, K: 0.5 mm; B, G, L, P: 0.2 mm; C, H, M, R: 0.1 mm.

### Eye field transcription factors are robustly expressed in the arrested OV of severely affected E10.5 *Porcn*^*CKO*^

Since OC morphogenesis is consistently and impaired the most, we proceeded with analyzing severely affected E10.5 *Porcn*^*CKO*^ (Figure 2F-J). The pan-ocular transcription factor PAX6 is expressed in the arrested *Porcn*^*CKO*^ OV and also present in the surface ectoderm (Figure 3B). The eye field transcription factor Six3 is required for neuroretinal specification the mammalian eye (Liu and Cvekl, 2017; Liu et al., 2010). SIX3 expression is not altered in the OV and surface ectoderm of *Porcn*^*CKO*^ (Figure 3D). LHX2 is not altered (Figure 3F), confirming that general ocular specification including the transition from eye field to OV and OV identity are maintained at later stages.

**Figure 3:**
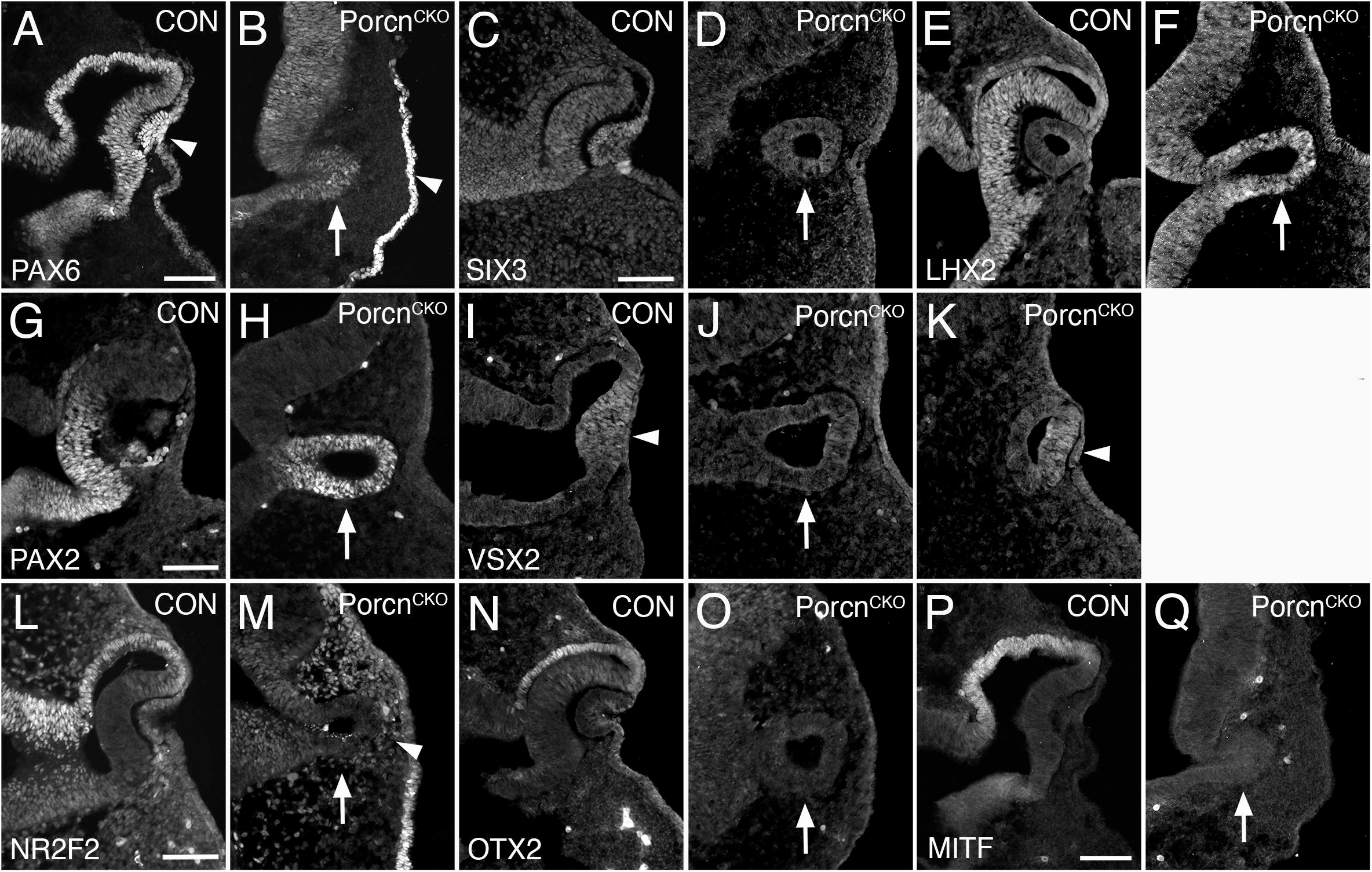
*Porcn* mutant embryos display defective regionalization at E10.5. Coronal view of control embryos (A, C, E, G, I, L, N, P) and *Porcn*^*CKO*^ embryos that were treated with tamoxifen at E7.4 (B, D, F, H, J, K, M, O, Q). (A) During normal eye development, PAX6 is expressed in ocular tissues and in the overlying surface ectoderm (arrowhead). (B) In the mutant OV, PAX6 expression is expressed (arrow) and is present in the surface ectoderm (arrowhead). (C, D) SIX3 expression is not altered in the OV and surface ectoderm of *Porcn*^*CKO*^ (D; arrow). (E, F) In control (E) and in *Porcn*^*CKO*^ (F; arrow) LHX2 is expressed throughout the OC and OV, respectively. (G) In control OC, PAX2 is restricted to the ventral OC. (H) In *Porcn*^*CKO*^, PAX2 expression is maintained throughout the OV (arrow). (I) In controls, the distal OC is tightly attached to the overlying lens ectoderm and expresses VSX2 (arrowhead). (J) In *Porcn*^*CKO*^ with loss of close contact, VSX2 is not detectable (arrow). (K) When distal OV and lens ectoderm are closely associated, some cells in the distal OV in *Porcn*^*CKO*^ can express VSX2 (arrowhead). (L) NR2F2 expression shown in control embryos. (M) In the OV of *Porcn*^*CKO*^, NR2F2 expression is missing (arrow). NRF2 expressing POM cells are abnormally present between distal OV and surface ectoderm (arrowhead). (N, O) OTX2 is normally robustly expressed in the presumptive RPE of the OC but is absent in the mutant OV (O; arrow). (P) MITF is an early differentiation marker restricted to the presumptive RPE. (Q) In the OV of *Porcn*^*CKO*^, MITF expression is not detectable (arrow). Scale bars A, C, G, L, P: 0.1 mm

### Porcn mutant embryos display defective regionalization of the arrested OV at E10.5

The paired homeobox transcription factor PAX2 is initially expressed throughout the OV and is later confined to the ventral OC and optic stalk (Burns et al., 2008; Nornes et al., 1990). In *Porcn*^*CKO*^, PAX2 expression is maintained throughout the OV (Figure 3H), consistent with failure of OC formation and dorsoventral regionalization. During normal eye development, proximodistal regionalization into RPE and retina occurs in the advanced OV (Fuhrmann, 2010; Fuhrmann et al., 2014; Miesfeld and Brown, 2019). In *Porcn*^*CKO*^ with loss of close contact due to POM accumulation between the distal OV and surface ectoderm (Figure 2), the strictly retina-specific CVC homeodomain transcription factor VSX2 is not expressed (Figure 3J) (Burmeister et al., 1996; Green et al., 2003). However, if close contact occurs in *Porcn*^*CKO*^, some cells in the presumptive retina can express VSX2 (Figure 3K), consistent with the requirement of lens-derived FGF for retina specification (Cai et al., 2013). Concerning proximal regionalization, we observed a complete loss of the key regulatory transcription factors NR2F2, OTX2 and MITF (Figure 3M, O, Q) that are required for RPE differentiation (Bumsted and Barnstable, 2000; Hodgkinson et al., 1993; Martinez-Morales et al., 2001; Nguyen and Arnheiter, 2000; Tang et al., 2010). NRF2 expression confirms abnormal abundance of POM between distal OV and lens ectoderm (Figure 3 M). Together, our results demonstrate that dorsoventral and proximodistal regionalization is impaired in the arrested OV of *Porcn*^*CKO*^ at the time when OC morphogenesis normally commences.

### Decrease of proliferation, survival and local NR2F2 and OTX2 expression in the *Porcn*^*CKO*^ OV at E9.5

To elucidate a potential mechanism underlying abnormal ocular growth obvious in E10.5 *Porcn*^*CKO*^, we analyzed embryos one day earlier, following tamoxifen induction at E7.4-E7.5. At E9.5, *Porcn*^*CKO*^ show slightly decreased telencephalic vesicles and mid-hindbrain regions (Figure 4B). LEF1 expression in the dorsal OV and adjacent dorsal forebrain neuroepithelium in *Porcn*^*CKO*^ embryos is unaltered, suggesting that Wnt/β −catenin signaling is intact at this age (Figure 4D). MITF expression is not affected in the presumptive RPE of *Porcn*^*CKO*^ (Figure 4F). However, other RPE markers NR2F2 and OTX2 start to be reduced dorsally in the distal OV domain (Figure 4H, J) suggesting that some aspects of regionalization can be affected at this age. We observed increased cell death in the *Porcn*^*CKO*^ OV (Suppl. Figure 2). To determine effects on proliferation, we quantified the number of E9.5 OV cells incorporating EdU (Figure 4K-O). In *Porcn*^*CKO*^, the number of EdU-labeled cells was significantly decreased in the entire OV (Figure 4L, M) that is specifically due to a significant decrease in the ventral OV by 27% (Figure O). Therefore, *Porcn* inactivation affects survival and proliferation of ocular progenitors in the ventral OV resulting in severe growth defects in *Porcn*^*CKO*^.

**Figure 4:**
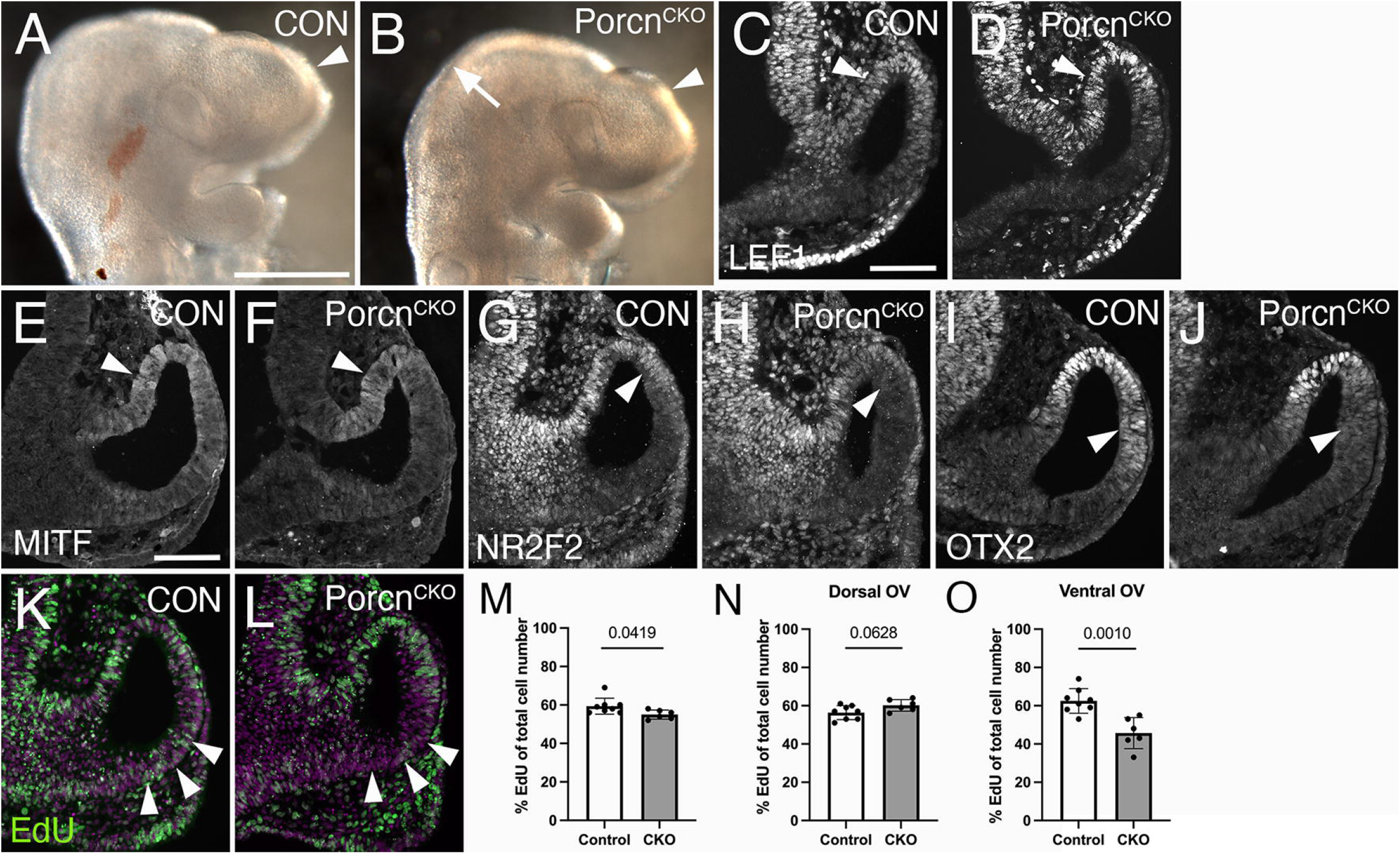
Decrease of proliferation, survival and local NR2F2 and OTX2 expression in the *Porcn*^*CKO*^ OV at E9.5. Coronal view of control embryos (A, C, E, G, I, K) and *Porcn*^*CKO*^ embryos, induced at E7.4-7.5 (B, D, F, H, J, L). (A) Telencephalic vesicle in E9.5 control embryo (arrowhead). (B) *Porcn*^*CKO*^ show slightly smaller telencephalic vesicles (arrowhead) and mid-hindbrain regions (arrow). (C) In controls, LEF1 is expressed in the dorsal OV (arrowhead). (D) In *Porcn*^*CKO*^ embryos, LEF1 is unaltered (arrowhead). (E, F) MITF expression appears unaffected in the presumptive RPE of *Porcn*^*CKO*^ (F; arrowhead). (G, H) In the mutant RPE, NR2F2 can be slightly downregulated in the dorsal OV (H; arrowhead). (I, J) OTX2 can start to be reduced dorsally in the distal OV domain in Porcn^CKO^ (J; arrowhead). (K, L) Edu incorporation (green) in control (K) and in Porcn^CKO^ OV (L), showing reduced number of EdU-labeled cells in the ventral OV (L; arrowheads). Magenta: Dapi. (M-O) Quantification of EdU labeled cells in the entire OV (M), dorsal OV (N) and ventral OV (O). In Porcn^CKO^, the number of EdU-labeled cells is significantly decreased in the entire OV (M) and in the ventral OV (O). Data are means ± s.d. Student’s T-test was applied for statistical analysis, and p-values are indicated on the horizontal lines in each graph (significance < 0.05). Scale bars A: 0.5 mm; C, E: 0.1 mm

## Discussion

Our results show that *Porcn* inactivation either before or during eye field formation leads to severely arrested OVs, likely due to an increase in cell death and downregulation of proliferation. In *Porcn*^*CKO*^, the RPE key regulatory transcription factors OTX2, MITF and NR2F2 are downregulated. POM accumulates between distal OV and adjacent surface ectoderm, preventing tight association, and the retina-specific gene VSX2 is not properly expressed. Arrested eye vesicles in *Porcn*^*CKO*^ do not invaginate, resulting in failed invagination of the OV into an OC. Our studies reveal a novel role for Porcn in the OV as the earliest obvious morphological stage of eye development and a continued requirement during OC morphogenesis, recapitulating severe microphthalmia in FHD.

### Effect of Porcn disruption on Wnt signaling

We first determined when potential effectors of Wnt signaling are affected by *Porcn* inactivation at E6.5. Wnt/β −catenin signaling is normally not active until OV invagination starts; the pathway readouts Axin2 and BATgal reporter are absent in the OV around E8.75 (8 somites) (Liu et al., 2010). Here, we observed that another readout, LEF1, starts to become weakly expressed around E9.0 in the dorsal OV of controls (13 somites; Figure 1C). In *Porcn*^*CKO*^ embryos, LEF1 is slightly downregulated in the dorsal OV (Figure 1D) suggesting commencement of a pathway response. Indeed, in the E9.5 OV, LEF1 is absent, demonstrating complete shutdown of the pathway, OV morphogenesis is abnormal (Figure 1M). Thus, it takes at least 2.5 days after tamoxifen treatment to detect initial effects of *Porcn* inactivation on Wnt/β −catenin signaling and approximately 3 days to observe defects in gene expression and eye morphogenesis.

Since Wnt/β-catenin signaling needs to be suppressed in the anterior neural plate, we reasoned that any requirement for Porcn would be likely due to a need for the non-canonical Wnt pathway. Inactivation of *Porcn* also prevents potentially confounding, concomitant upregulation of Wnt/β-catenin signaling. Consistent with this, arrested growth of the OV has not been observed by early Wnt/β −catenin pathway inactivation in other studies (Hagglund et al., 2013). We observed that the putative non-canonical Wnt pathway readout pJUN is normally not robustly expressed in the E9.5 OV (not shown), and F-actin shows normal apicobasal localization in *Porcn*^*CKO*^ (Figure 1). However, we cannot exclude effects on other potential non-canonical Wnt targets in *Porcn*^*CKO*^. Furthermore, our strategy of *Porcn* inactivation may be not quick enough to affect EFTF expression in the eye field between E7.5 and E8.5 (see above).

However, it may be challenging to perform *Porcn* inactivation before E6.5 without causing most severe developmental defects allowing to determine whether Porcn promotes EFTF expression and eye field formation.

### Porcn is not required for maintenance of EFTF expression in the OV and OC

Eye field formation with expression of EFTFs in mouse starts around E7.5. To determine Porcn’s role during the earliest stages of eye morphogenesis, we induced inactivation at E6.5. Analysis of E8.75-9.5 *Porcn*^*CKO*^ embryos revealed that expression of the EFTFs LHX2 and OTX2 is unaffected at E9.0, and that LHX2 is still present 3 days after tamoxifen induction (Figure 1F, H, Q). This also applies to *Porcn*^*CKO*^ induced at E7.5 and analyzed at E10.5; the EFTFs PAX6, LHX2 and SIX3 are expressed (Figure 3B, D, F). Therefore, our data demonstrates that Porcn is not required to maintain EFTF expression in the OV.

### Porcn is required for initiation of RPE differentiation in the OV

Our results are consistent with previous studies showing that *Porcn* inactivation prevents maintenance of RPE differentiation in the OC, most likely due to loss of Wnt/β −catenin signaling. RPE-specific inactivation of the pathway effector β −catenin in the early OC interferes with further RPE differentiation (Fujimura et al., 2009; Hagglund et al., 2013; Westenskow et al., 2009). Loss of RPE fate is likely caused by a failure to transactivate RPE gene expression, therefore, the mutant RPE transdifferentiates into retina (Fujimura et al., 2009; Westenskow et al., 2009). Here we show that RPE differentiation fails to initiate in *Porcn*^*CKO*^ embryos induced at E6.5, since the early RPE marker OTX2 is absent in the dorsal E9.5 OV (Figure 1S). The POM is essential for RPE differentiation and a source for Wnt ligands, in addition to ocular tissues (Bankhead et al., 2015; Bassett et al., 2010; Carpenter et al., 2015; Fuhrmann et al., 2000; Gage et al., 1999). Thus, ubiquitous depletion of Porcn may interfere with RPE differentiation cell and non-cell autonomously. To examine a cell autonomous requirement, we attempted to disrupt *Porcn* using a more restricted *Cre* line, *Hes*^*CreERT2*^ (Kopinke et al., 2011; Yun et al., 2009). We administered up to 0.12 mg/g tamoxifen around E6.8 and observed no obvious phenotype in *Porcn* mutant embryos (n=4; not shown). We detected mosaic RosaR26 reporter expression in only one E10.5 embryo (n=3; not shown) suggesting that *Hes*^*CreERT2*^ may not be sufficiently activated at this time point. However, we showed in an earlier study that *Porcn* disruption causes consistent RPE differentiation defects only when performed simultaneously in ocular and extraocular tissues in the OV (Bankhead et al., 2015). We propose that Wnts are also redundantly produced and available within the eye field and adjacent tissues.

### Porcn is required for ocular growth by promoting proliferation and survival of ocular progenitors

Our study demonstrates that *Porcn* inactivation results in severe morphogenesis defects 3 days later, either at the OV (E9.5) or OC stage (E10.5). The *Porcn*^*CKO*^ OV or OC is small and OV expansion or OC invagination fails, respectively. Wnt ligands can directly regulate proliferation and survival of ocular progenitors (Burns et al., 2008; Hagglund et al., 2013). Analysis of cell death (TUNEL) and proliferation (EdU incorporation) showed robustly downregulated survival of ocular progenitors and proliferation in the ventral OV (Suppl. Figure 2; Figure 4K-O). Significantly reduced proliferation during OC morphogenesis has been observed in embryos with germ line mutation of early ocular regulatory genes, for example Pax6^sey/sey^, Lhx2, BMP7, Hes1 and Mab21L2 (Grindley et al., 1995; Lee et al., 2005; Morcillo et al., 2006; Porter et al., 1997; Yamada et al., 2004; Yun et al., 2009). However, Pax6 and Lhx2 are expressed in the OV of *Porcn*^*CKO*^, and Pax6, BMP7, Hes1 and Mab21L2 mutants do not show a robust morphogenesis defect until E10.5. Therefore, to our knowledge, the ocular phenotype in *Porcn*^*CKO*^ is unique because it shows an earlier requirement for proliferation and survival of ocular progenitors.

### Porcn inactivation around the eye field stage recapitulates microphthalmia and anophthalmia in FDH patients

*Porcn*^*CKO*^ display severe microphthalmia 3 days after tamoxifen treatment before or during the eye field stage. *Porcn*^*CKO*^ embryos induced at E6.5 and harvested at E11 showed severe developmental abnormalities not feasible for further analysis at later time points (not shown). Lhx2 mutants exhibit a similar OV morphogenesis defect leading to anophthalmia subsequently (Porter et al., 1997). Thus, we hypothesize that *Porcn* inactivation before the OV stage ultimately results in anophthalmia, consistent for an early role of Porcn in FDH. A case report for 18 FDH patients revealed a high incidence of ophthalmologic abnormalities (77%), including microphthalmia (44%) and anophthalmia (11%) (Gisseman and Herce, 2016). We propose that *Porcn* inactivation around the eye field stage represents a novel mouse model recapitulating severe microphthalmia and anophthalmia in humans.

## Conclusion

Using temporally controlled, conditional inactivation in mouse, our studies reveal a novel role for *Porcn* in regulating growth and morphogenesis of the optic vesicle and optic cup, via a requirement in proliferation, survival, and regionalization of ocular gene expression, recapitulating severe microphthalmia in Focal Dermal Hypoplasia.

## Ethics statement

Animal procedures were reviewed and approved by the Institutional Animal Care and Use Committee at Vanderbilt University Medical Center.

## Author contributions

SF designed experiments, supervised the research, carried out experiments presented in the manuscript, analyzed the data, and wrote the manuscript. SR carried out experiments, analyzed data and edited the manuscript. MMA carried out experiments, analyzed data and edited the manuscript. CC carried out experiments, analyzed data and edited the manuscript.

## Acknowledgements

Special thanks to Ethan Lee and Yukio Saijoh for expert advice. Many thanks to Maria Elias and Katrina Hofstetter for excellent technical support. We thank Ed Levine and his team for helpful comments and discussions. Special thanks to Charles Murtaugh (University of Utah, Salt Lake City, UT) for kindly providing the conditional *Porcn* allele and the *Cre* line *Hes1*^*CreERT2*^.

## Funding

This work was supported by the National Institutes of Health (R01 EY024373 to S.F., T32 HD007502Training Grant in Stem Cell and Regenerative Developmental Biology to S.R., Core Grants P30 EY008126, EY14800); a Catalyst Award to S.F. from Research to Prevent Blindness Inc./American Macular Degeneration Foundation, an unrestricted award to the Department of Ophthalmology and Visual Sciences from Research to Prevent Blindness, Inc.; the Vanderbilt University Medical Center Cell Imaging Shared Resource Core Facility (Clinical and Translational Science Award Grant UL1 RR024975 from National Center for Research Resources).

## Legends

**Supplemental Figure 1: RosaR26 reporter activation**.

Coronal view of E9.0 embryonic heads, induced with tamoxifen at E6.5 (A, B) and E9.5 embryos (C, D), induced at E7.4. (A, B, D) *Porcn*^*CHET*^ and *Porcn*^*CKO*^ embryos show widespread expression of β-galactosidase protein, in contrast to controls without *Cre* (C; m FL). Arrows point to OVs in each image.

Scale bar: 0.1 mm

**Supplemental Figure 2: Detection of apoptotic cell death by TUNEL**.

(A-C) Coronal view of representative images of TUNEL-labeled E9.5 sections induced with tamoxifen at E7.4. Compared to *Cre*-negative controls (A; m FL), overall TUNEL signal appears increased throughout the tissues in *Porcn*^*CHET*^ and *Porcn*^*CKO*^ embryos (B, C). In the ventral *Porcn*^*CKO*^ OV, more TUNEL-labeled cells are detectable (C), compared to *Porcn*^*CHET*^ (B)

Arrowheads mark the extent of TUNEL-labeled region in the ventral OV (B, C). Scale bar: 0.1 mm

**Suppl. Table 1:**
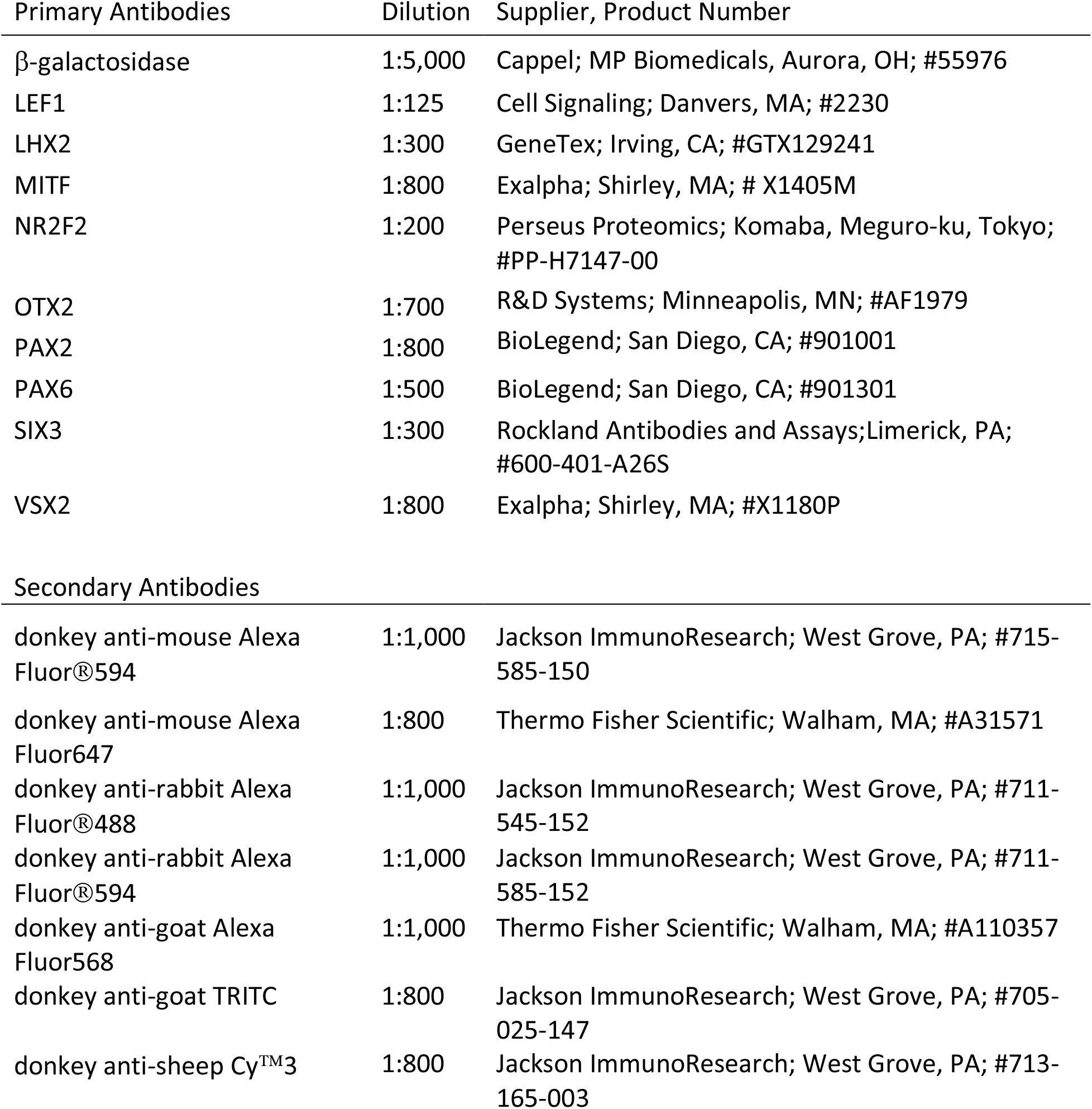
Antibodies used in this study.

## Notes

### Competing Interest Statement

The authors have declared no competing interest.

