## Supplementary figures and images for "Porcn is essential for growth and invagination of the mammalian optic cup"

### Suppl Figure 1

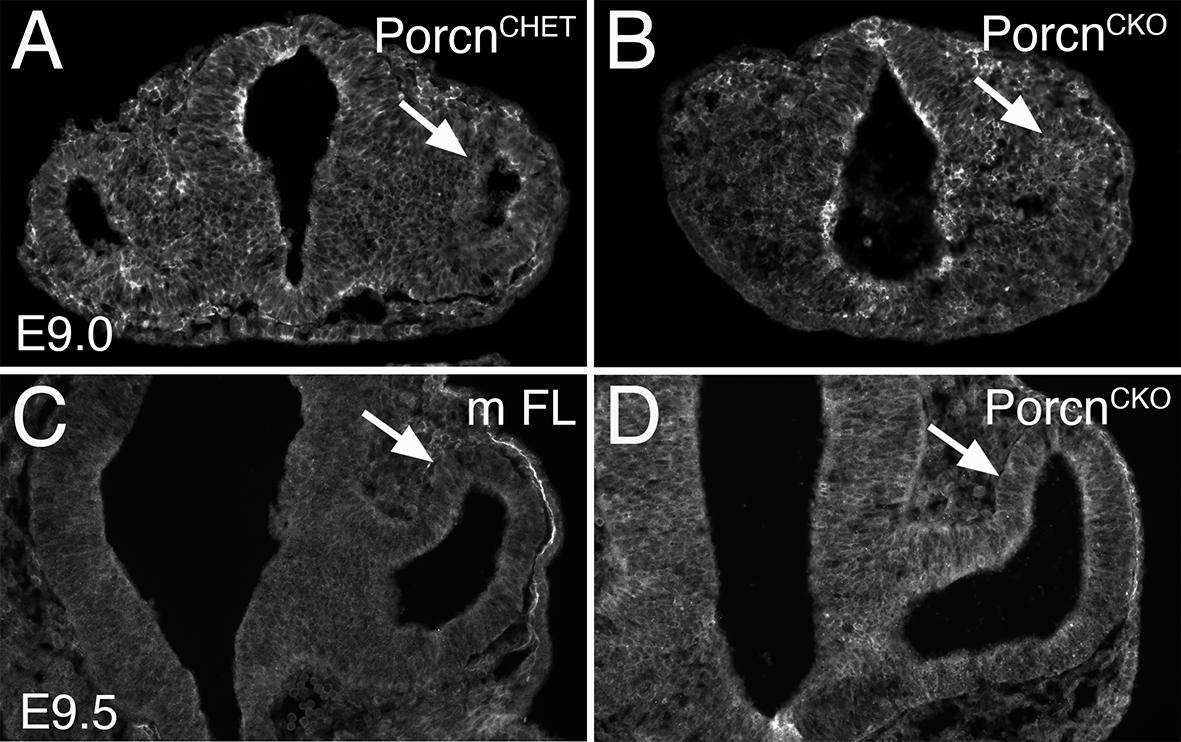

### Suppl Figure 2

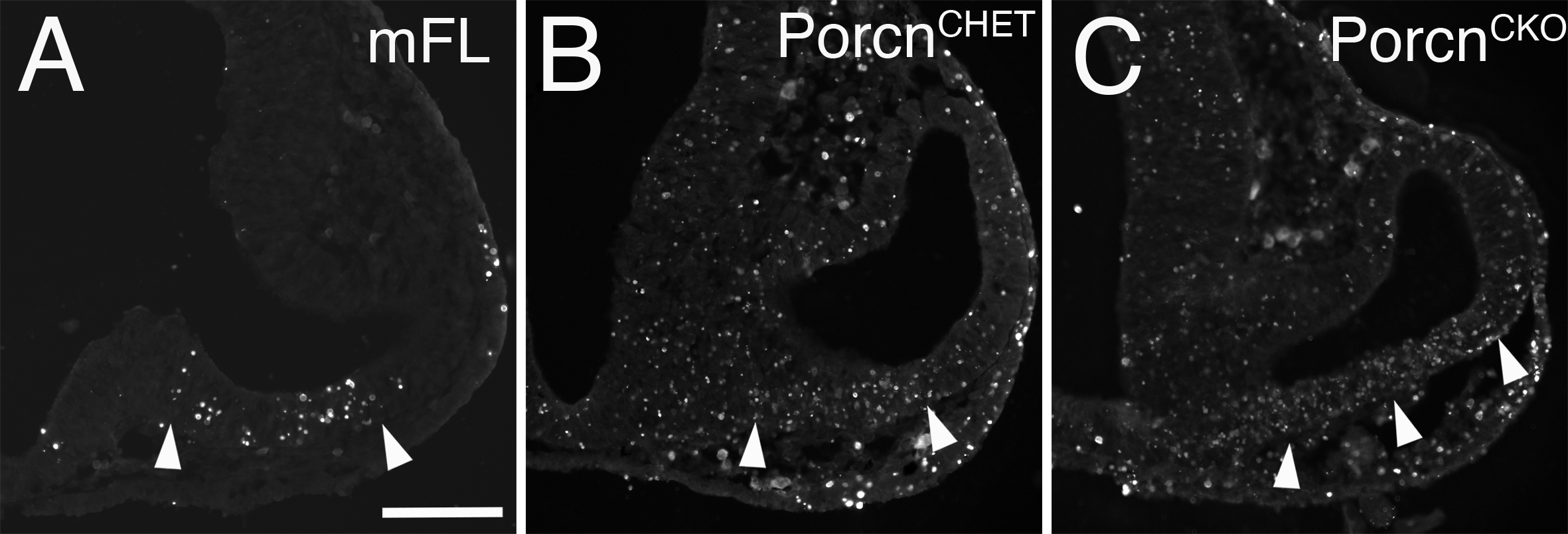
